# Discovery and characterisation of new phage targeting uropathogenic *Escherichia coli*

**DOI:** 10.1101/2024.01.12.575291

**Authors:** Shahla Asgharzadeh Kangachar, Dominic Y. Logel, Ellina Trofimova, Hannah X Zhu, Julian Zaugg, Mark A. Schembri, Karen D. Weynberg, Paul R. Jaschke

## Abstract

Antimicrobial resistance (AMR) is increasing at an escalating rate with few new therapeutic options in the pipeline. Urinary tract infections (UTIs) are one of the most prevalent bacterial infections globally and are particularly prone to becoming recurrent and antibiotic resistant. The aim of this study was to discover and characterise new bacterial viruses (phage) against uropathogenic *Escherichia coli* (UPEC), which is the leading cause of UTIs. Six phages from the *Autographiviridae* family and *Guernseyvirinae* sub-family were isolated from wastewater and sequenced. The length of the isolated phage genomes was between 39,471 bp and 45,233 bp, with a GC content between 45.0% and 51.0%, and 57 to 84 predicted coding sequences (CDS) per genome. These phages were found to infect between 25 – 75% of the twelve UPEC strains tested. Using sequence comparison and predicted structural alignments, we show a similarity between the C-terminal domain of the tail fiber proteins of two phage that correlates with their host range. *In vitro* characterisation of phage cocktails against a single bacterial strain did not perform better than the best-performing phage, but did show synergistic improvement against a mixed UPEC strain population. Lastly, we measured the effectiveness of treatment with phage with different lytic kinetics in a sequential treatment and found it was improved over single phage treatment.

## INTRODUCTION

Bacterial infectious diseases were the leading cause of human mortality prior to antibiotic discovery, and their use caused an immediate reduction in mortality rates in subsequent years (Aminov 2017). However, bacterial strains developed resistance to antibiotics in a short time, and in the case of penicillin, only two years after its discovery (Walsh 2000). Now, in the post-antibiotic era, antimicrobial resistance (AMR) has increased to such an extent that it has been declared a worldwide crisis by the World Health Organisation (Ventola 2015). By 2050, the mortality rate attributed to antimicrobial resistance (AMR) is projected to escalate to 10 million annually, with one in five human bacterial infections in Australia, North America and Europe predicted to be antibiotic resistant (O’Neill 2014; Sugden et al. 2016; Stirrups 2019).

The rate of newly approved antibacterial agents for clinical usage, including antibiotics, has not been able to address the escalating occurrence of antibiotic-resistant bacteria (Ribeiro da Cunha et al. 2019). Phage therapy is a promising alternative to antibiotics, where bacterial viruses or bacteriophages (known as phages) are used to kill bacteria (Lemire et al. 2018), and much progress is being made in the effective use of phages in the clinic (Dedrick et al. 2019; Gordillo Altamirano and Barr 2019; Dedrick et al. 2023).

Urinary tract infections (UTIs) are one of the most prevalent bacterial infections, with ~400 million infections estimated globally in 2019 (Yang et al. 2022). UTIs disproportionately affect females, older adults, and people with diabetes mellitus or polycystic kidney disease (Suárez Fernández et al. 2021). The leading cause of UTIs is uropathogenic *Escherichia coli* (UPEC), with 80% of cases due to this causative group (Delcaru et al. 2016; Galtier et al. 2016; NTK Nhu 2023). Further, UTIs from UPEC can progress to more serious infections with the bacteria migrating into the blood via the bladder and renal system, eventually causing sepsis (Flores-Mireles et al. 2015). In bacterial UTIs, biofilm formation provides protection for slow-growing or dormant strains, increasing antibiotic resistance that in turn contributes to chronic and recurrent infection (Ballash et al. 2022). Recurrent UTIs occur in 20-30% of cases, and are linked to increased antibiotic usage and treatment failure.

Antibiotic use for uncomplicated UTIs can vary based on country and its underlying bacterial antibiotic resistance profile (Kang et al. 2018), but nearly always involves an empirical treatment pattern starting with broad-spectrum first-line antibiotics such as trimethoprim/sulfamethoxazole or nitrofurantoin. Lack of response from first-line treatment and in multidrug resistant infections will prompt treatment with antibiotics such as third-generation cephalosporins.

Using phages against UPEC in UTIs is an emerging treatment option due to their ability to reduce or eliminate antibiotic resistant bacterial populations that cannot be addressed with antibiotics alone (Zalewska-Piatek and Piatek 2020; Al-Anany et al. 2023). Phages can also be engineered for higher efficacy through release of antimicrobial agents at the site of infection (Du et al. 2023); they can be used synergistically with antibiotics (TorresLBarceló et al. 2016), and as a cocktail to eradicate the antibiotic resistant strains (Galtier et al. 2016; Esmael et al. 2021). Using phage cocktails has been reported to successfully treat bacterial infection (Kim et al. 2021; Alexyuk et al. 2022; Benala et al. 2023) though standardised methods are yet to be established (Kim et al. 2021; Alexyuk et al. 2022; Benala et al. 2023).

In this study our aim was to isolate and characterise novel phage against UPEC. Here we report the isolation, sequencing, host range characterisation, and determination of optimal *in vitro* cocktail compositions of six new bacteriophage against UPEC.

## MATERIALS AND METHODS

### Bacterial strains and culture condition

The UPEC bacterial strains used in this study are shown in Table 1. All bacterial cultures were grown in Lysogeny Broth Lennox (LB) (Sigma-Aldrich, 5 g/L NaCl, 10 g/L Tryptone, 5 g/L Yeast Extract, Product # L3022). For phage assays, cultures were grown to early exponential phase (OD_600_=0.3). Phage lytic performance in liquid culture was evaluated using 96 well plates (Costar, REF# 3599) with a total volume (phage and bacterial culture) of 200 µL in each well and monitored for a period of 24 hours at 37 °C using a CLARIOstar^®^ Plus multi-mode microplate reader (BMG Labtech, Offenburg, Germany). A multiplicity of infection (MOI) of 10 was used unless indicated otherwise.

**Table 1.** UPEC strains used in this study.

| Bacterial Strain | Serotype | Sequence type | LPS Outer Core Type | Isolated Phage | Host Reference |
| --- | --- | --- | --- | --- | --- |
| PA45B* | O2:H7:K1 | ST95 | R1 | vB_EcoM_SHAK7704 | (Goh et al. 2017) |
| S96EC* | O25b:H4:K5<sup>#</sup> | ST131 | K-12 | vB_EcoM_SHAK7854 | (Petty et al. 2014) |
| S112EC* | O25b:H4:K+ | ST131 | K-12 | vB_EcoM_SHAK7858 | (Petty et al. 2014) |
| MS7163* | O45:H7:K1 | ST95 | R1 | vB_EcoM_SHAK7163 | (Nhu et al. 2018) |
| S39EC* | O25b:H4 | ST131 | K-12 | vB_EcoM_SHAK9454 | (Petty et al. 2014; Phan et al. 2015) |
| CFT073 | O6:H1:K2 | ST73 | R1 | - | (Mobley et al. 1990) |
| PA7B* | O6:H1:K23 | ST73 | R1 | vB_EcoM_SHAK7693 | (Nichols et al. 2016) |
| S101EC* | O25b:H4:K5<sup>#</sup> | ST131 | K-12 | - | (Petty et al. 2014) |
| S79EC | O25b:H4:K1<sup>#</sup> | ST131 | K-12 | - | (Petty et al. 2014) |
| S129EC | O25b:H4 | ST131 | K-12 | - | (Petty et al. 2014) |
| S116EC | O25b:H4 | ST131 | K-12 | - | (Petty et al. 2014) |
| HVM2044 | O25b:H4:K5<sup>#</sup> | ST131 | K-12 | - | (Petty et al. 2014) |

\*Bacterial strains used for phage isolation.
#*In silico* capsule (K) prediction.

### Initial phage isolation and enrichment

Bacteriophage were isolated from sewage collected from the Luggage Point Wastewater Treatment Plant, Brisbane, Australia. The sludge suspension was allowed to settle at 4 °C overnight. The sludge supernatant was then dispersed into 50 mL Falcon tubes and centrifuged at 5,000 ×*g* for 10 minutes at 4 °C. The resulting supernatant was vacuum filtered using 0.22 µm pore size filters (Millipore, USA) to remove any bacteria and other contaminants. The filtrate was filtered again with a fresh 0.22 µm filter to ensure any contamination had been removed and was stored at 4 °C in the dark.

For bacteriophage enrichment, a single bacterial colony (listed in Table 1) was inoculated in 10 mL LB broth and incubated for 4 - 6 hours at 37 °C, 150 RPM. This culture was then split 1/500 into fresh LB broth and incubated at 37 °C, 150 RPM overnight. In a 250 mL flask, 100 µL of overnight bacterial culture was added to 30 mL double-strength LB broth supplemented with 2 mM CaCl_2_ and 2 mM MgCl_2_. To this starter culture, 20 mL of double-filtered sewage was added. Flasks were incubated at 37 °C, 50 RPM (gentle shaking) for 24 hours. The overnight incubated samples were centrifugated at 11,000 ×g for 12 minutes at 4 °C, followed by filtering the supernatant with 0.22 µm pore size filters to obtain the enriched phage lysate.

The presence of phage in the lysate was confirmed by performing a double-layer agar plate method plaque assay (Kropinski et al. 2009). Briefly, 100 µL of enriched phage lysate and 300 µL of bacterial host starter culture, were added to 3 mL LB agarose medium (0.4% agarose, 0.7% LB broth) and poured onto a 25 mL, already solidified agar plate and incubated at 37 °C overnight. Single plaques formed on the bacterial lawn were picked and isolated using a sterile pipette tip and stored in 1 mL SM buffer (8 mM MgSO_4_.7H_2_O; 50 mM Tris-Cl; 100 mM NaCl; pH 7.2). To ensure the purity of the isolated phage, three more sequential rounds of plaque assay were performed from the SM buffer containing a single isolated plaque. The isolated phages were tested for chloroform sensitivity and were stored in a LoBind Eppendorf tube (Cat# 0030108051) at 4 °C in the dark for further downstream analysis.

### Initial phage propagation

An initial phage propagation was performed to increase the volume of phage lysate to proceed with further phage purification. To do so, a single bacterial colony was inoculated in 10 mL of LB broth in a 50 mL falcon tube and incubated 4-6 hours at 37 °C, 150 RPM; 200 µL of this liquid culture was then added to 10 mL fresh LB broth and incubated overnight at 37 °C, 150 RPM. A mixture of 1 mL overnight bacterial culture and 10 mL double-strength LB broth supplemented with 2 mM CaCl_2_ and 2 mM MgCl_2_ was prepared. Tubes were incubated at 37 °C, 150 RPM for 2.5 hours, followed by adding 100 µL of isolated phage in SM buffer. Samples were incubated overnight at 37 °C, 150 RPM. The overnight incubated samples were centrifugated at 4,000 ×g for 20 minutes at 4 °C, followed by passage through a 0.22 µm syringe filter. A spot assay (described below) was performed to assess the presence of successful phage amplification.

### Phage purification via Ultracentrifugation

OptiPrep^TM^ (Sigma-Aldrich) density gradient ultracentrifugation (Beckman Coulter Optima XPN-100 Ultracentrifuge, SW 41 Ti rotor) was performed for each isolated sample. The density gradient tube included 3 mL 50%, 2.5 mL 40%, 2.5 mL 30% and 2 mL 20% Opti Prep^TM^ from bottom to top, and the sample was loaded as the last fraction on the top. Samples were centrifuged at 288,000 ×*g* for 18 hours at 4 °C. The bands were collected with syringes and stored at 4 °C in the dark for further downstream analysis.

Anion exchange chromatography (GE Healthcare PD MidiTrap™ G-25) was used to remove Optiprep from purified phage, using the spin protocol as per the manufacturer’s instructions.

To increase phage titre following ultracentrifugation, two rounds of subsequent propagation were performed as described above. Following small-scale amplification, a larger-scale propagation was performed using a final volume of 1-litre culture for each sample (5 flasks of 200 mL for each sample).

The phage lysate was concentrated using either Centricon (Centricon® Plus-70 Centrifugal Filter Units, Merck) or Amicon (Amicon® Ultra-15 Centrifugal Filter Unit, Merck) filter units with centrifugation (3,200 ×g, 4 °C, variable times based on each individual sample).

### Bacteriophage titering

The concentration of each isolated phage was identified by performing a double-layer agar plate method plaque assay as described above. The plaque assay was performed in three technical replicates, and the plates, including between 10 and 100 plaques, were chosen for plaque count.

### Bacteriophage host range analysis via spot assay

The spectrum of the host range for the isolated phages was determined using the spot lysis assay as follows. 300 µL of bacterial host starter culture was added to 3 mL LB agarose medium (0.4% agarose, 0.7% agar) and poured into an LB agar plate and let to be solidified. 10 µL of the purified phage lysate was spotted on top of the solidified agarose. The droplets were allowed to air dry for approximately 30 minutes at room temperature in a biosafety cabinet and incubated at 37 °C overnight. The result was interpreted based on plaque formation classified into three different categories: +++ for the efficiency of plating (EOP) of 80% and above, ++ for EOP between 40% and 79%, and + for the EOP between 20% and 39%.

### Transmission electron microscopy

To visualise the morphology of isolated phages, transmission electron microscopy was performed. Purified and concentrated phage lysate was used as the sample, and the 200-mesh carbon-coated copper grids (ProSciTech code: GSCU150CC) were negatively stained as follows: 14 µL of the 10^11^ cfu/mL of sample was dropped on the grid for 5 minutes, followed by two washes with Milli-Q water. Then 14 µL of uranyl acetate (UA) 2% was dropped on the grid and incubated for one minute at room temperature, and grids were air dried in the dark. Grids were examined, and images were taken using the transmission electron microscope JEM-1011 (JEOL^TM^, Tokyo, Japan), with an accelerating voltage of 80 kV. ImageJ (v1.53e (Abràmoff et al. 2004)) was used to add scale bars to the images.

### Phage genomic DNA extraction

Phage genomic DNA was extracted from concentrated phage lysate using a phage DNA isolation kit (Norgen Biotek Corp., Thorold, Canada) according to the manufacturer’s protocol, with the exception that molecular biology grade water was used for elution. All optional steps were conducted. The extracted DNA was then cleaned up and concentrated using the Genomic DNA Clean & Concentrator (Zymo Research Corp.) kit.

The quality and quantity of the extracted phage DNA were determined by TapeStation (Agilent Technologies 4200 TapeStation) using Genomic DNA ScreenTape. The samples for TapeStation analysis were prepared as follows: 10 µL of genomic DNA sample buffer was added to 1 µL of sample or ladder and mixed with shaking for 60 seconds at 2,000 RPM, followed by a brief spin.

### Genome sequencing

Both Illumina short read and Oxford Nanopore Technology (ONT) (Oxford Nanopore Technologies, United Kingdom) long-read sequencing platforms were employed for whole-genome sequencing. For short-read sequencing, the samples were submitted to the Australian Centre for Ecogenomics (ACE). The library preparation method was Nextera DNA Flex, and NextSeq (Illumina) genome sequencing instrument was used to sequence the samples. For long-read sequencing, the MinION Nanopore sequencer was used. The SQK-RBK004 Rapid Barcoding Kit was used to prepare the library and was run on the MinION flow cell. Library preparation and sequencing were performed according to the manufacturer’s protocol (RBK_9054_v2_revP_14Aug2019 updated on 09/12/2020) with all optional steps included and an 18-hour run time.

### Genome assembly and annotation

Reads generated from sequencing were assembled through the following workflow. MinION FAST5 data was basecalled with guppy 4, and filtered using Nanofilt (De Coster et al. 2018) (v 2.7.0, length=1000, quality=9, headcrop=30). Long reads passing this quality-control step were then assembled using Flye (Kolmogorov et al. 2019) (v2.9, flye --meta). Assemblies were then polished using Racon (Vaser et al. 2017) (v1.4.20, racon_match=8, racon_mismatch=-6, racon_gap=-8, racon_window-length=500) for three rounds, followed by Medaka (v 1.4.4; https://github.com/nanoporetech/medaka). Finally, Illumina short-reads were trimmed using Trimmomatic (Bolger et al. 2014) (v0.39) and used to polish the assemblies with Pilon (Walker et al. 2014) (v1.24).

Structural and functional annotation of the assembled genomes were performed using the CPT Galaxy platform (Ramsey et al. 2020; The Galaxy 2022) (https://cpt.tamu.edu/galaxy-pub). Briefly, for the structural annotation, three different tools were used, including Glimmer (Delcher et al. 2007), MetaGene annotator (Noguchi et al. 2008), and Sixpack (Rice et al. 2000). Independently from the Galaxy platform, assemblies were annotated with Pharokka (Bouras et al. 2022) (v1.4.1) including the integrated MASH (Fast genome and metagenome distance estimation using MinHash) analysis (Ondov et al. 2016). Then manually, the overlaps and disagreements between different tools were reviewed, and either the longest predicted CDS, or the CDS that was annotated to be a member of a Prokaryotic Virus Remote Homologous Group (PHROG) (Terzian et al. 2021) was retained in the final annotation.

A phylogenetic tree of phage large terminase sequences was constructed using VipTree (Nishimura et al. 2017) v1.9.1 (https://www.genome.jp/viptree/) based on the proteomic tree concept original outlined by Rohwer and Edwards(Rohwer and Edwards 2002). VipTree was used with the following parameters: dsDNA, host categories of reference viruses from prokaryote, auto gene prediction with procaryotic genetic code.

The presence of virulence factor and antibiotic resistance genes, including both chromosomal point mutations and acquired AMR genes within the assembled phage genomes, were identified using VirulenceFinder v 2.0 (Joensen et al. 2014; M Tetzschner et al. 2020), and ResFinder v4.1 (Camacho et al. 2009; Zankari et al. 2017; Bortolaia et al. 2020), respectively. The threshold for minimum percent identity and minimum length were set to 90% and 60%, respectively. PhaBOX and PhaTYP were used, with default parameters, to determine phage lifestyles (Shang et al. 2022; Shang et al. 2023).

Annotated phage genome sequences were compared to other phage genomes using clinker (Gilchrist and Chooi 2021) hosted by the online CompArative GEne Cluster Analysis Toolbox (CAGECAT) (https://cagecat.bioinformatics.nl/).

The GenBank accession numbers for the phage identified in this work are:

vB_EcoM_SHAK7163 (OR594179), vB_EcoM_SHAK7854 (OR594182),

vB_EcoM_SHAK7704 (OR594181), vB_EcoM_SHAK9454 (OR594184),

vB_EcoM_SHAK7693 (OR594180), vB_EcoM_SHAK7858 (OR594183).

### Detection of antiphage defence systems in bacterial genomes

We used the web version of PADLOC (Payne et al. 2022) (accessed 02/2023), DefenseFinder (Tesson et al. 2022) (accessed 03/2023), and CRISPRDetect (Biswas et al. 2016) (accessed 03/2023), to identify antiphage defence systems present in UPEC host genomes. GenBank accession numbers used: DAAZYG000000000.1 (S112EC), DAAZXO000000000.1 (S129EC), DAAZZJ000000000.1 (S39EC), DAAZYH000000000.1 (S101EC), DAAZWZ000000000.1 (S79EC), DAAZYU000000000.1 (S96EC), DAAZVR000000000.1 (HVM2044), DAAZXW000000000.1 (S116EC), NZ_CP051263.1 (CFT073), NZ_CP026853.1 (MS7163), DADZYB000000000.1 (PA7B), NZ_CP021288.1 (PA45B). Default parameters in all tools were used and subtypes of systems in “other” type (e.g., DMS_other) were excluded from analysis.

### Outer core LPS typing

Bacterial outer core types were manually classified based on the previously established *E. coli* outer core LPS operon structures (Heinrichs et al. 1998) and LPS genes were identified from those operons in each strain using the UniProt (Consortium 2022) Blast service (https://www.uniprot.org/blast).

### Protein structural prediction

Protein sequences from the phage CDSs were analysed through the publicly available AlphaFold (Jumper et al. 2021; Varadi et al. 2022) software (colab.research.google.com/github/deepmind/alphafold/blob/main/notebooks/AlphaFold.ipynb). Sequences were searched individually using default options. All models were visualised through PyMOL v2.3.4 (Schrödinger, LLC). To determine structural similarities between the predicted protein models all protein sequences were aligned using the PyMOL align function with default parameters.

## RESULTS

In the current study, our goal was to isolate and characterize naturally occurring phages against UPEC strains, identify genes associated with phage host range, and investigate different combinations of the phage against these UPEC strains.

### Bacteriophage isolation and host range determination

To isolate host-specific bacteriophages, we obtained wastewater from the Luggage Point Wastewater Treatment Plant, Brisbane, Australia. We isolated six bacteriophages that produced plaques on a range of UPEC strains (Table 1). Each isolated bacteriophage was named based on the International Committee on Taxonomy of Viruses (ICTV) guidelines (Adriaenssens and Brister 2017) (Table 1).

To evaluate the host range of each isolated phage, we performed a spot lysis test of each phage against 12 UPEC strains (Table 2). All phages infected at least one strain beyond the strain used to isolate it. vB_EcoM_SHAK7858 displayed the broadest host-range, producing plaques on nine out of 12 hosts, closely followed by vB_EcoM_SHAK7704, vB_EcoM_SHAK9454, and vB_EcoM_SHAK7693 which all produced plaques on eight strains (Table 2). S112EC is the most phage-susceptible UPEC strain, and is permissive for all six isolated phages. The second most phage-susceptible UPEC strains are S116EC and S79EC, and are infected by five out of six isolated phages.

**Table 2.**
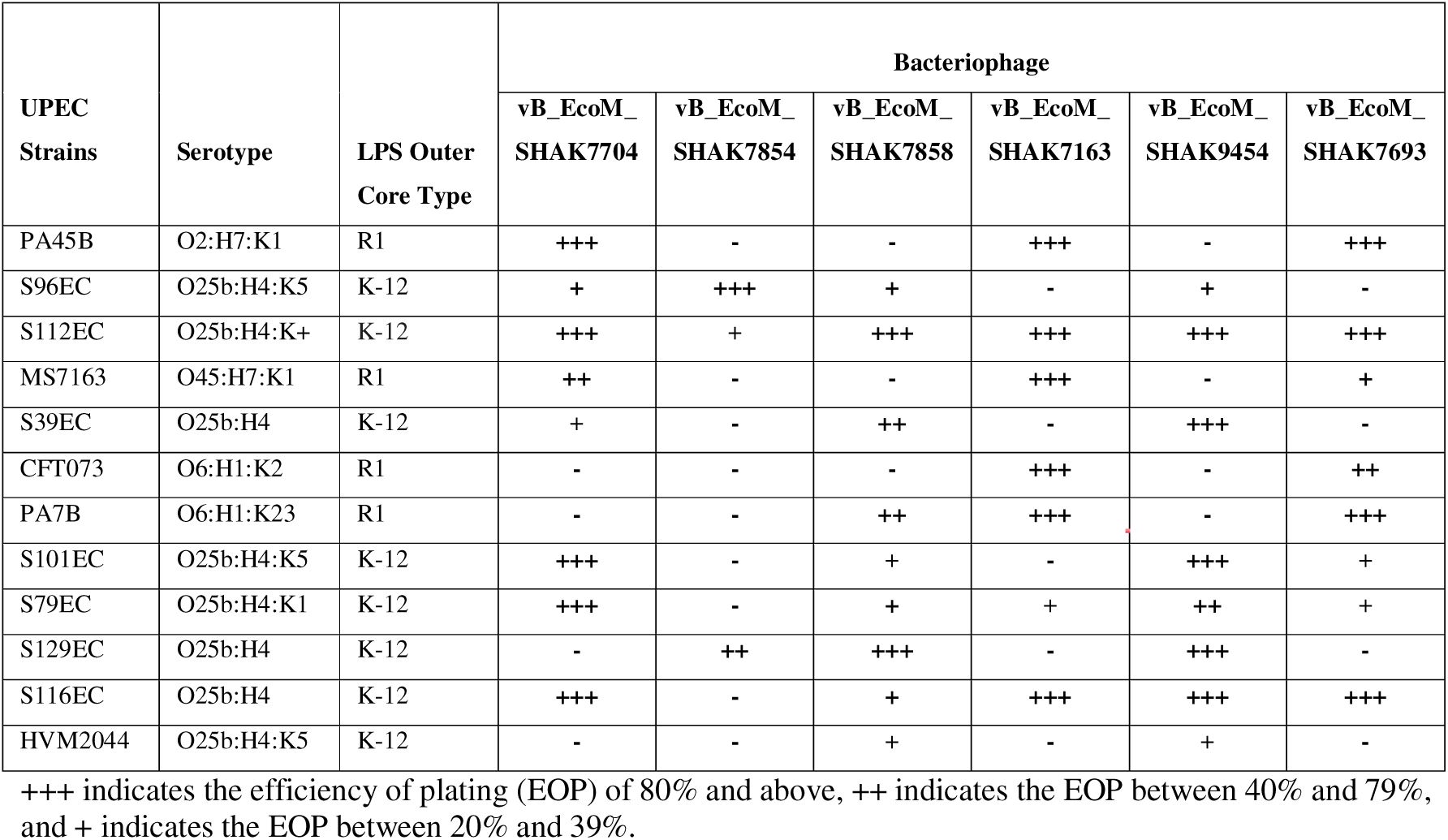
Bacterial host range analysis for isolated phages against UPEC strains.

| UPEC Strains | Serotype | LPS Outer Core Type | Bacteriophage |  |  |  |  |  |
| --- | --- | --- | --- | --- | --- | --- | --- | --- |
|  |  |  | vB_EcoM_SHAK7704 | vB_EcoM_SHAK7854 | vB_EcoM_SHAK7858 | vB_EcoM_SHAK7163 | vB_EcoM_SHAK9454 | vB_EcoM_SHAK7693 |
| PA45B | O2:H7:K1 | R1 | +++ | - | - | +++ | - | +++ |
| S96EC | O25b:H4:K5 | K-12 | + | +++ | + | - | + | - |
| S112EC | O25b:H4:K+ | K-12 | +++ | + | +++ | +++ | +++ | +++ |
| MS7163 | O45:H7:K1 | R1 | ++ | - | - | +++ | - | + |
| S39EC | O25b:H4 | K-12 | + | - | ++ | - | +++ | - |
| CFT073 | O6:H1:K2 | R1 | - | - | - | +++ | - | ++ |
| PA7B | O6:H1:K23 | R1 | - | - | ++ | +++ | - | +++ |
| S101EC | O25b:H4:K5 | K-12 | +++ | - | + | - | +++ | + |
| S79EC | O25b:H4:K1 | K-12 | +++ | - | + | + | ++ | + |
| S129EC | O25b:H4 | K-12 | - | ++ | +++ | - | +++ | - |
| S116EC | O25b:H4 | K-12 | +++ | - | + | +++ | +++ | +++ |
| HVM2044 | O25b:H4:K5 | K-12 | - | - | + | - | + | - |

+++ indicates the efficiency of plating (EOP) of 80% and above, ++ indicates the EOP between 40% and 79%, and + indicates the EOP between 20% and 39%.

Transmission electron microscopy was performed next to visualize the phage capsid morphologies (Figure 1). We observed four phages with a *Podoviridae/Autographiviridae* capsid type (vB_EcoM_SHAK7163, 7704, 7854, and 9454), and two phages with a *Siphoviridae* capsid type (vB_EcoM_SHAK7693 and 7858) (Figure 1).

**Figure 1.**
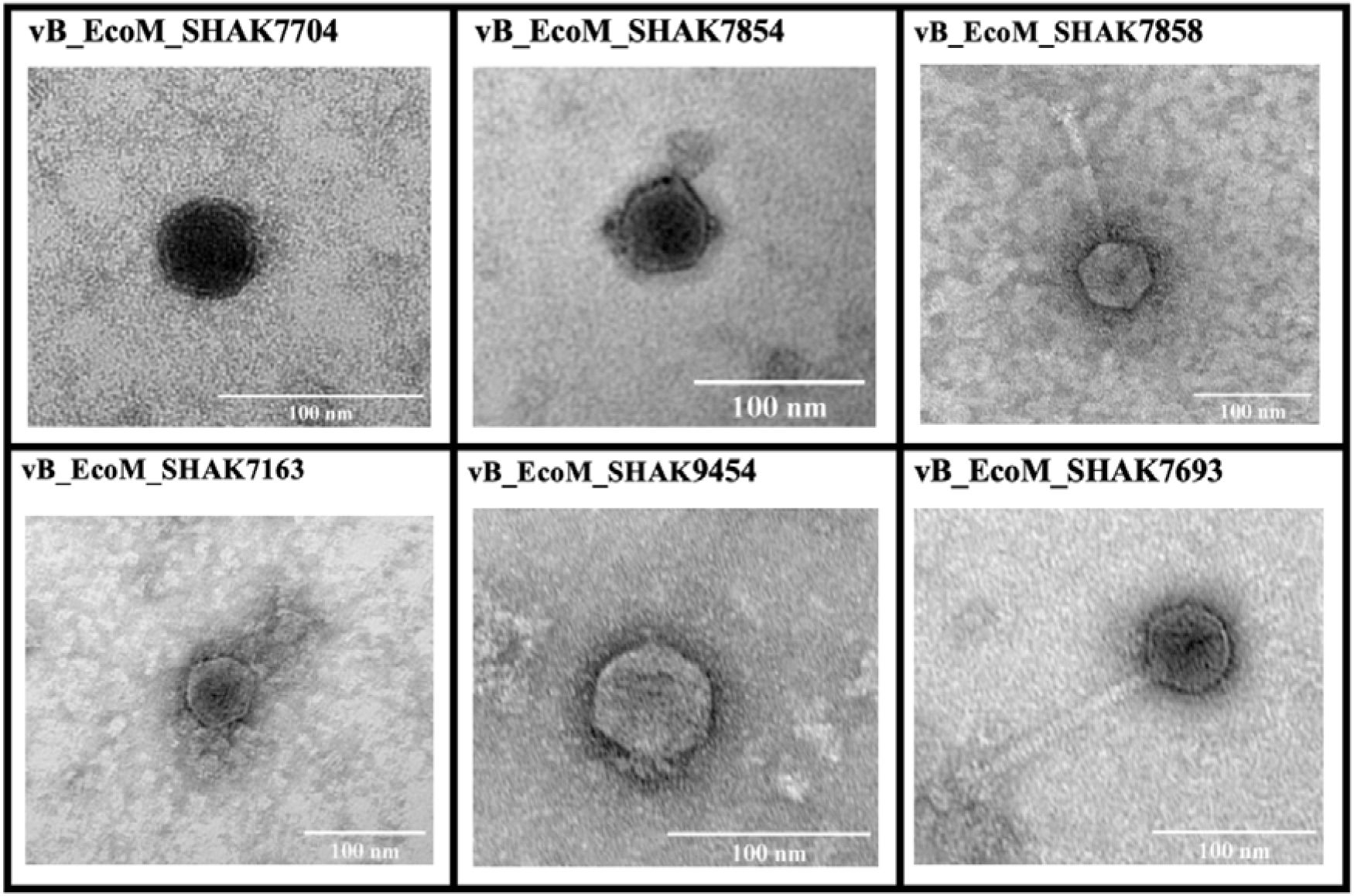
The morphology of isolated phages visualised with TEM. Images taken using JEM-1011 with an accelerating voltage of 80 kV.

### Bacteriophage genome sequencing and annotation

To more precisely classify and characterise the isolated phages, we sequenced their genomes via both short-read and long-read sequencing methods. The phage genomes varied in length, ranging between 39,471 bp to 45,233 bp and in GC content between 45.0% and 51.0% (Table 3).

**Table 3.** Phage genome sequence and annotation characteristics.

| Phage | Length<br>(bp) | GC<br>content<br>(%) | Predicted<br>CDSs | CDS<br>predictions<br>assigned to<br>a PHROG<br>(% of CDS<br>predictions) | Closest phage<br>match | Assigned<br>Genus |
| --- | --- | --- | --- | --- | --- | --- |
| vB_EcoM_SHAK7163 | 44,656 | 45.0 | 64 | 55 (85.9%) | Escherichia<br>phage K1E | *Vectrevirus* |
| vB_EcoM_SHAK7704 | 44,446 | 45.0 | 64 | 56 (87.5%) | Escherichia<br>phage K1E | *Vectrevirus* |
| vB_EcoM_SHAK7854 | 39,734 | 50.0 | 57 | 50 (87.7%) | Escherichia<br>phage<br>vB_EcoP-PM139 | *Kayfunavirus* |
| vB_EcoM_SHAK9454 | 39,471 | 50.0 | 57 | 50 (87.7%) | Escherichia<br>phage HC13 | *Kayfunavirus* |
| vB_EcoM_SHAK7693 | 45,233 | 51.0 | 84 | 68 (81.0%) | Escherichia<br>phage<br>vB_EcoS_fPo<br>Eco01 | *Kagunavirus* |
| vB_EcoM_SHAK7858 | 44,517 | 51.0 | 83 | 72 (86.7%) | Escherichia<br>phage<br>vB_EcoS-22664BS2 | *Kagunavirus* |

MASH (Fast genome and metagenome distance estimation using MinHash) analysis (Ondov et al. 2016) showed the nearest genome matches in the INPHARED database (Cook et al. 2021) were all from phages with *E. coli* hosts (Table 3). Using these matches we were able to classify our isolated phages at the genus level (Table 3) which was confirmed by a large terminase subunit phylogenetic tree (Figure S1). These results also corresponded well with our morphological categorisation by transmission electron microscopy (Figure 1).

Four of the isolated phage were assigned to the *Autographiviridae*. Phages vB_EcoM_SHAK7163 and vB_EcoM_SHAK7704 within the *Autographiviridae* were assigned to the *Vectrevirus* genus within the *Molineuxvirinae* sub-family, while vB_EcoM_SHAK7854 and vB_EcoM_SHAK9454 were assigned to the *Kayfunavirus* genus within the *Studiervirinae* sub-family. By contrast, vB_EcoM_SHAK7693 and vB_EcoM_SHAK7858 were assigned to *Kagunavirus* within the *Guernseyvirinae* sub-family.

Structural and functional annotation for the six isolated phages was performed via CPT Galaxy functional workflow (Ramsey et al. 2020) and Pharokka (Bouras et al. 2022), with manual curation. Each genome was predicted to contain between 57 and 84 protein coding sequences (CDSs) (Table 3), with 81.0 - 87.7% of CDS annotations successfully assigned to a prokaryotic virus remote homologous group (PHROG) (Terzian et al. 2021). ResFinder and VirulenceFinder did not identify any acquired antibiotic resistance or acquired virulence genes. Similarly, tRNAscan-SE and Aragorn did not predict any tRNA or tmRNA genes in any of the phage genomes. Unsurprisingly, each protein function category occurred in a similar proportion within each of the three phage sub-families (Figure 2).

**Figure 2.**
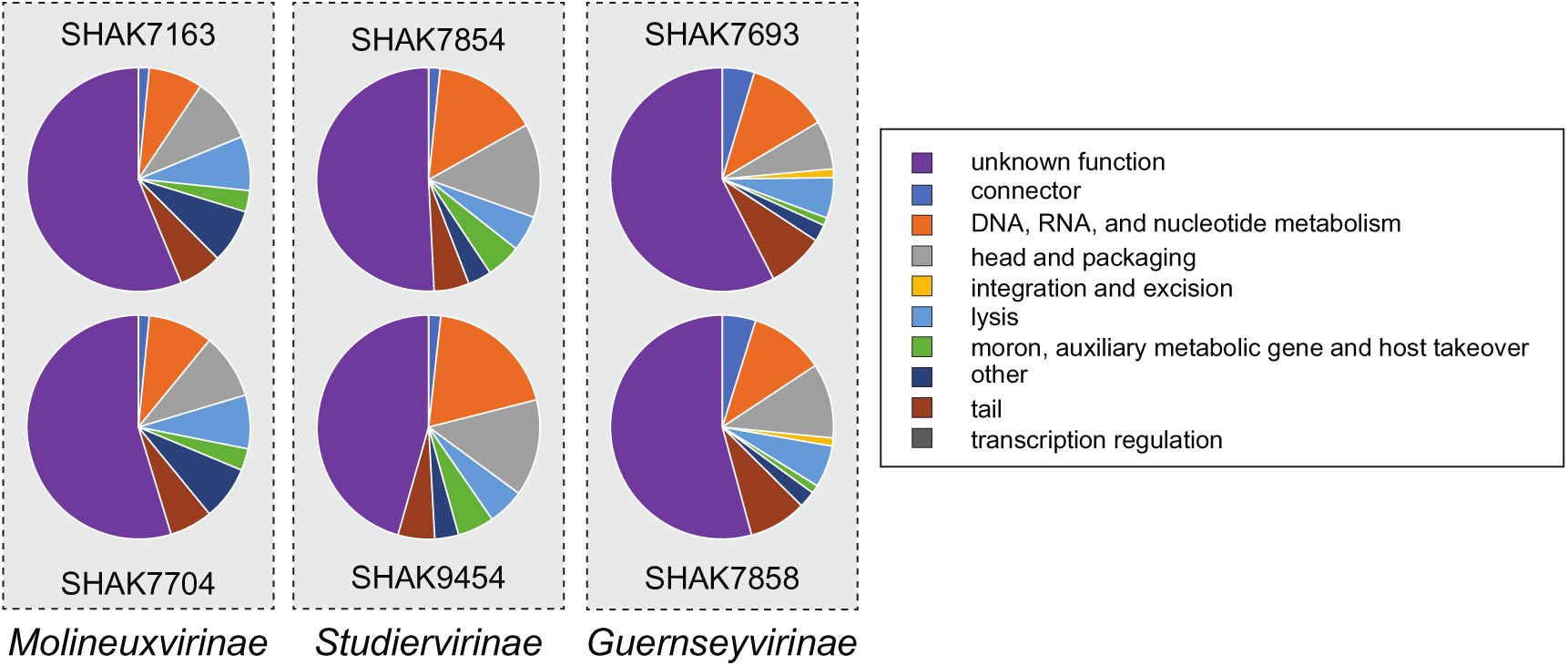
Protein CDS category breakdown across the isolated phages. Annotation categories assigned using PHANOTATE and PHROGs.

Pairwise comparison of the isolated phage genomes within each of the three sub-families showed a high degree of synteny and CDS sequence similarity (Figure 3). Comparison of each isolated phage with its closest identified sequence match also shows a high degree of synteny and sequence similarity (Figures S2 and S3). We also compared the four isolated phages within the *Autographiviridae* family with the most well understood member of the family, T7 phage. This comparison showed that all genomes display some degree of synteny, but that *Molineuxvirinae* sub-family members vB_EcoM_SHAK7163 and 7704 share very few homologous CDS with T7, while the *Studiervirinae* sub-family members vB_EcoM_SHAK7854 and 9454 are syntenic and share a significant number of homologous CDS with T7 (Figure S4).

**Figure 3.**
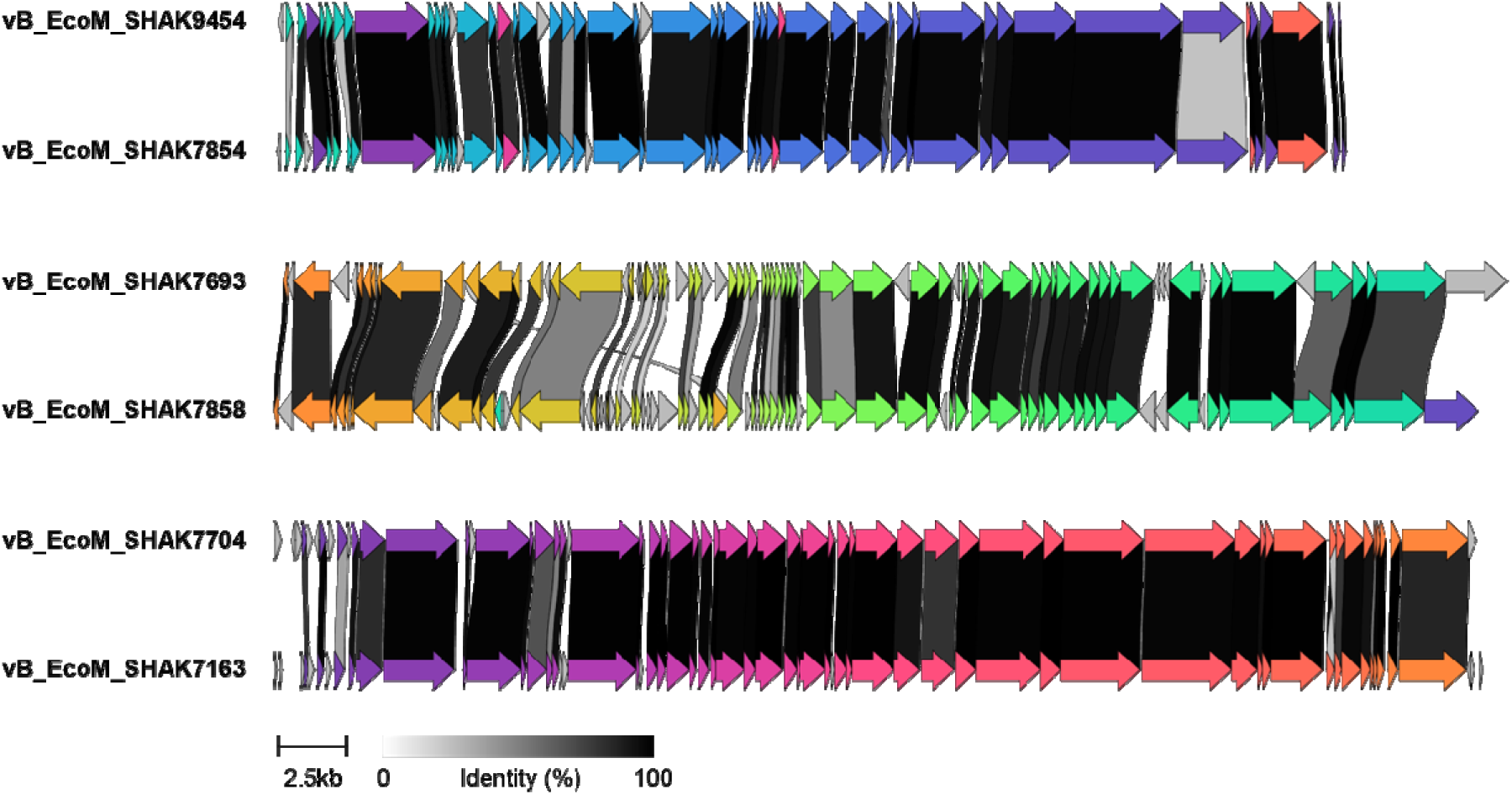
Genome CDS comparison of the isolated phages using clinker. Homologous CDSs are in the same colour and linked through grayscale bars based on percentage amino acid identity.

Interestingly, vB_EcoM_SHAK7693 and 7858 contain one CDS annotated in the integration and excision category. Looking closer, SHAK7693_0058 and SHAK7858_0053 CDS are annotated as “excisionase and transcriptional regulator” products. The CDSs are categorised into PHROG number 66 with a top hit to gene product NC_023692_p35 of Mycobacterium phage BigNuz. Inspection of the individual members of this PHROG did not reveal strong experimental support for this product function or anything that would indicate it is indicative of a lysogenic lifestyle. Similarly, PhaBOX analysis showed very high probability of a lytic lifestyle for both vB_EcoM_SHAK7693 and vB_EcoM_SHAK7858 using PhaTYP.

The only phage counter-defence proteins identified are in vB_EcoM_SHAK7163, 7854, and 7704 genomes which all contain an “ocr-like anti-restriction” protein product. All the proteins are highly similar to T7 protein gp0.3 (Ocr) which is a B-DNA mimic that protects phage DNA from restriction and modification by type I DNA restriction/modification enzymes (Atanasiu et al. 2002; Consortium 2022).

To understand the antiphage defence capabilities of the UPEC strains used in this work we analysed their genome sequences using the Prokaryotic Antiviral Defence LOCator (PADLOC), DefenseFinder, and CRISPRDetect bioinformatics tools (Biswas et al. 2016; Payne et al. 2022; Tesson et al. 2022). The results showed a range of 10 – 15 defence systems per strain, with type I, II, and IV restriction-modification systems present in nearly all genomes (Table S1). Other common phage defence systems identified in these strains included PsyrTA, AbiE, Lamassu, Mok Hok Sok, Mokosh type II, retrons, and CBASS type I (Bernheim and Sorek 2020; Tesson et al. 2022; Georjon and Bernheim 2023).

### Host range and tail fiber or spike proteins

Isolated phage genomes were compared to determine if there were any detectable correlations between host range and CDSs. Phage vB_EcoM_SHAK9454 (*Kayfunavirus*) and vB_EcoM_SHAK7858 (*Kagunavirus*) were two of the isolated phage with the broadest host ranges, and they were compared with vB_EcoM_SHAK7854 (*Kayfunavirus*) which had the narrowest host range (Table 2).

We focused our analysis on the tail fiber or spike protein from each phage, which is associated with host interaction. Global sequence alignment of the tail fiber or spike proteins between the three phages showed that the N-terminal regions of vB_EcoM_SHAK7854 and vB_EcoM_SHAK9454 were highly similar over the first 26.6% and 30.9% of the proteins, respectively (Figure 4A). By contrast, vB_EcoM_SHAK7858 and vB_EcoM_SHAK9454 shared extensive similarity between their sequences outside of the N-terminal portion of the proteins, covering 77.2% and 70.2% of the total protein length, respectively (Figure 4B). AlphaFold structural prediction of these three proteins showed a similar pattern between the pairwise comparisons and structural conservation between vB_EcoM_SHAK7854 and vB_EcoM_SHAK9454 in the N-terminal domain, vB_EcoM_SHAK7858 and vB_EcoM_SHAK9454 in the central and C-terminal domains, and little detectable structural similarity between vB_EcoM_SHAK7854 and vB_EcoM_SHAK7858 at either N- or C-terminus (Figure 4C and Figure S5).

**Figure 4.**
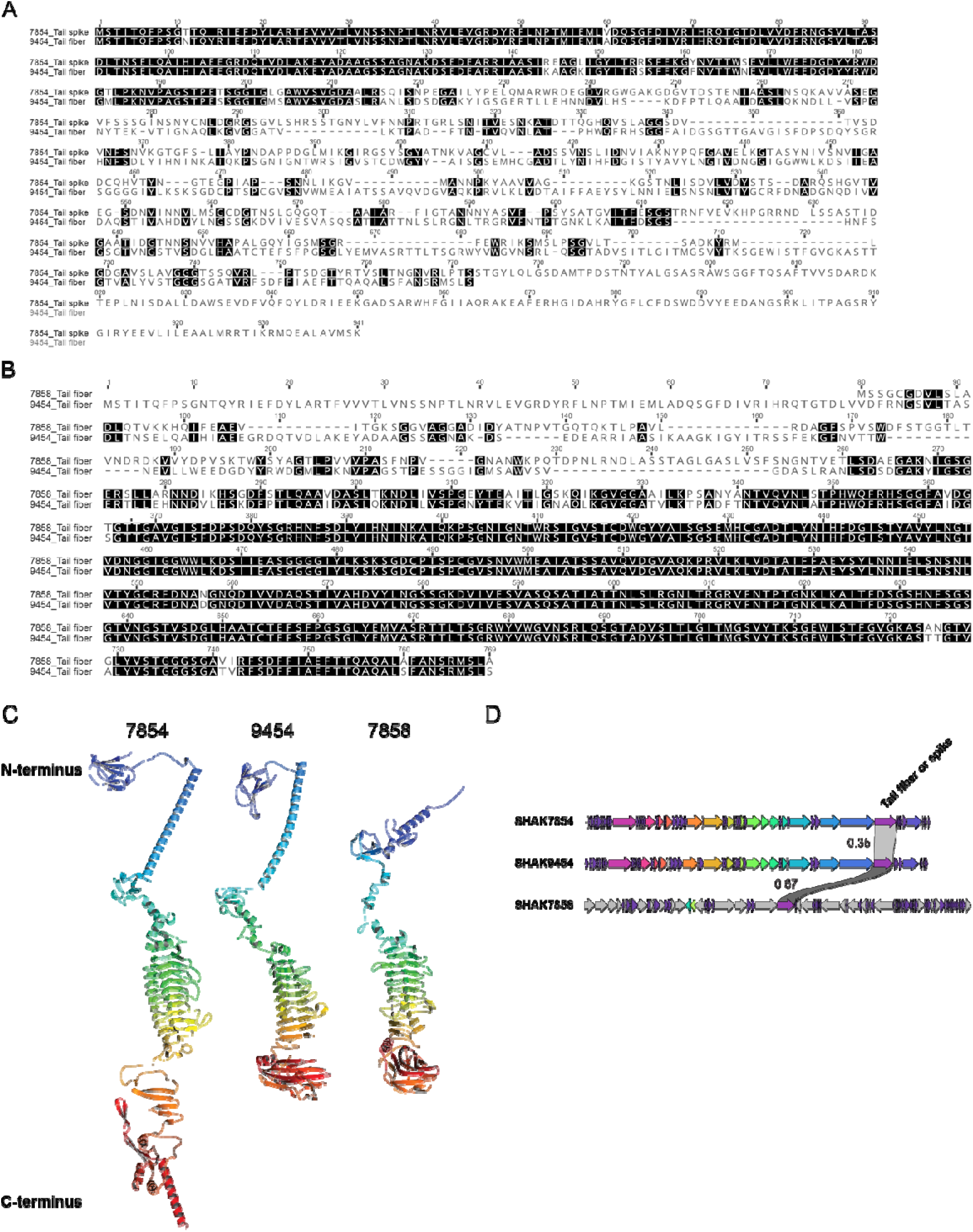
Tail fiber or spike protein C-terminus sequence and predicted structure correlates with host range characteristics. (A) Pairwise global sequence alignment between vB_EcoM_SHAK7854 tail spike protein and vB_EcoM_SHAK9454 tail fiber protein. (B) Pairwise global sequence alignment between vB_EcoM_SHAK7858 and vB_EcoM_SHAK9454 tail fiber proteins. (C) AlphaFold structural predictions of the tail fiber or spike proteins. Protein ribbons coloured in rainbow from N-terminus (blue) to C-terminus (red). (D) Tail fiber or spike gene location in phage genomes. Global protein sequence pairwise identity fraction shown.

This analysis showed the two *Kayfunavirus* phage (vB_EcoM_SHAK7854 and vB_EcoM_SHAK9454) shared N-terminal sequence and structure where the tail fiber or spike connects to the capsid, while the *Kagunavirus* vB_EcoM_SHAK7858 had a markedly different N-terminus (Figure 4C). By contrast, the C-terminus is known to be the point where the tail fiber or spike binds to the host receptor (Nobrega et al. 2018).

### Determination of phage *in vitro* bacterial killing efficiency

To determine the efficiency of the isolated phages against UPEC strains we tested both single phage and rational mixtures of phage. Based on our phage host range analysis (Table 2), which showed efficient lysis of PA45B by both vB_EcoM_SHAK7163 and vB_EcoM_SHAK7693, we tested these phages against this UPEC strain separately and as a cocktail. Phage vB_EcoM_SHAK7693 reduced culture absorbance sharply below the uninfected control after two hours, and this was maintained with slow recovery over the 24 hours (Figure 5). By contrast, phage vB_EcoM_SHAK7163 rapidly reduced culture absorbance after one hour, but culture density quickly recovered over the next six hours. A cocktail of these two phages shows changes to culture absorbance that closely matches that of vB_EcoM_SHAK7163 alone (Figure 5).

**Figure 5.**
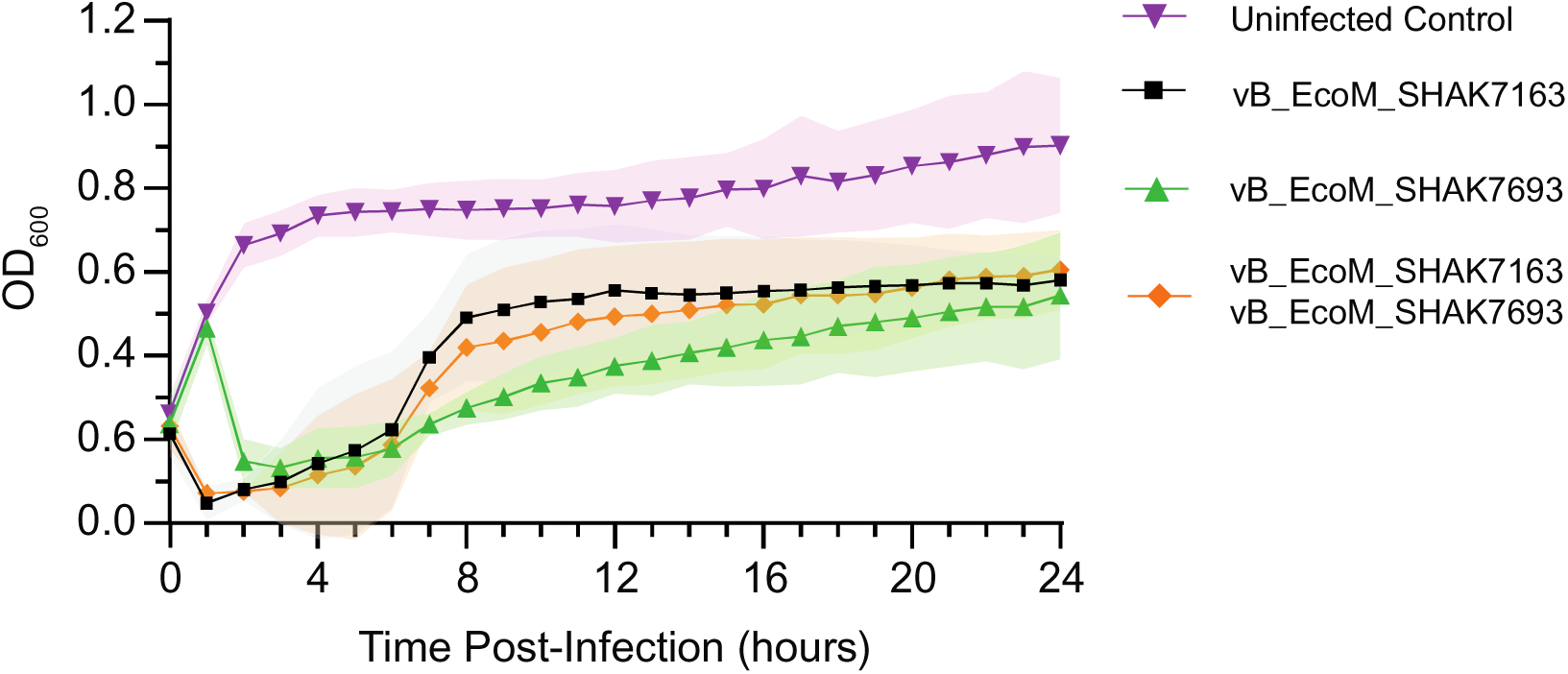
Performance of phage vB_EcoM_SHAK7163 and vB_EcoM_SHAK7693 alone and in combination against UPEC strain PA45B.

Because of our observations that phage cocktail infection patterns were driven exclusively by the effect of a single member of the mixture, our next experiment tested the effect of phage cocktails against mixtures of two bacterial hosts. Cocktails of two, three, and four phages (Table S2) were applied to 1:1 mixtures of bacterial strains S96EC and PA45B and were followed for 24 hours. The mixtures contained phage that were able to infect at least one of the bacterial strains used.

Of the 11 conditions tested, there was one cocktail that showed a synergistic effect between the two phages used (Figure 6). In the first eight hours after phage introduction, the combination of vB_EcoM_SHAK7854 and vB_EcoM_SHAK7693, reduced culture absorbance significantly better (p value < 0.01) compared to either single phage treatment (Figure 6).

**Figure 6.**
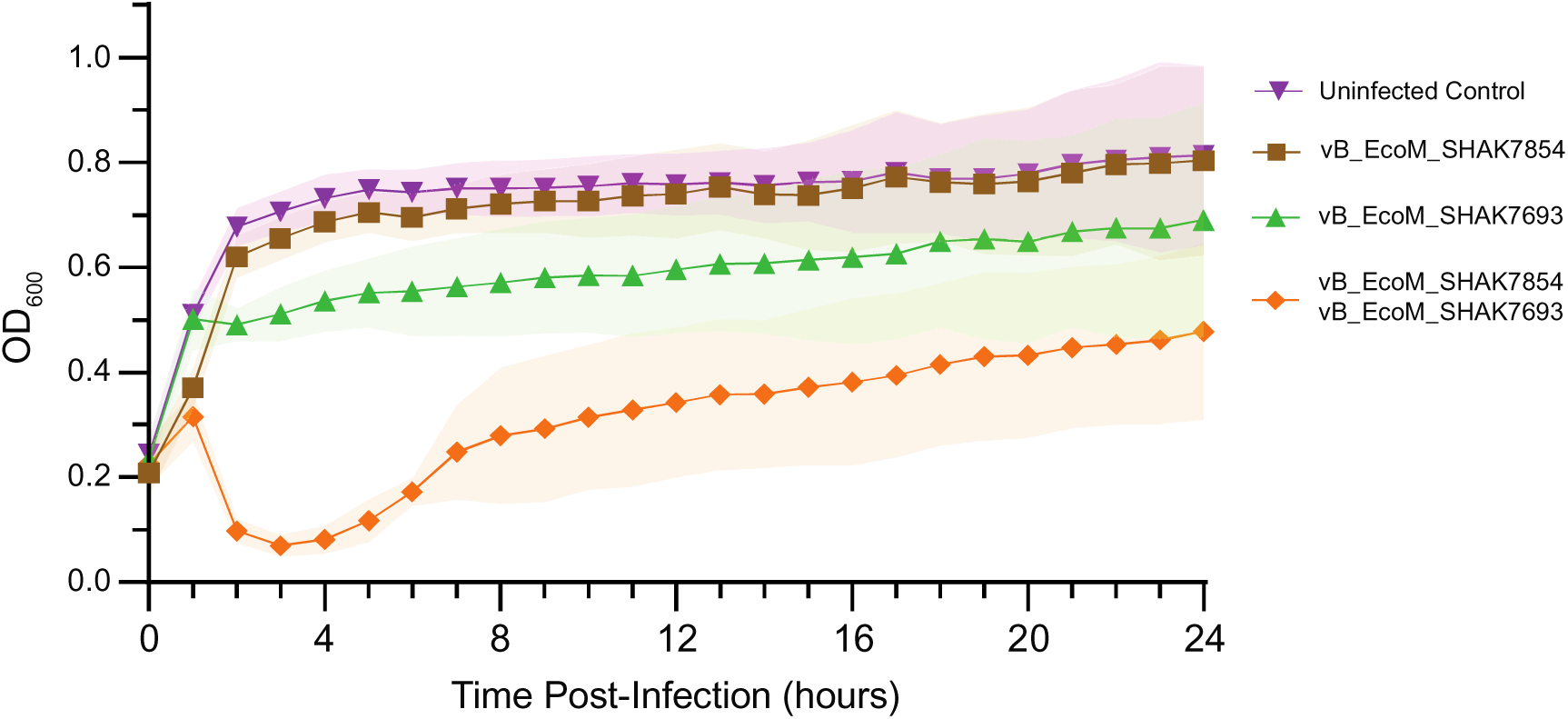
A phage cocktail of vB_EcoM_SHAK7854 and vB_EcoM_SHAK7693 against a 1:1 mixture of S96EC and PA45B shows synergistic activity.

### Sequential treatment with two different phages more effectively suppresses bacterial population than pooled initial dosing

To determine the effect that multiple phage treatments had on suppressing bacterial growth compared to only one initial phage application, we performed an experiment using sequential phage treatments. The S112EC strain was selected for this experiment because we had isolated one phage (vB_EcoM_SHAK7858) against this strain which showed quick initial suppression of bacterial growth, while a different phage (vB_EcoM_SHAK7693) had low initial suppression but a more long-term suppression effect (Figure 7). We measured 21 combinations of phage, varying dosing time and MOI (Table S3). Initial infection with vB_EcoM_SHAK7858 followed by sequential treatment with vB_EcoM_SHAK7693 at two, four, six, and 12 hours post-infection suppressed culture density at 16 - 24 hours significantly (p value <0.05) more than a single treatment of vB_EcoM_SHAK7858 alone (Figure 7). This dual treatment seemed to effectively combine the quick suppression of vB_EcoM_SHAK7858 with the more durable suppression of vB_EcoM_SHAK7693.

**Figure 7.**
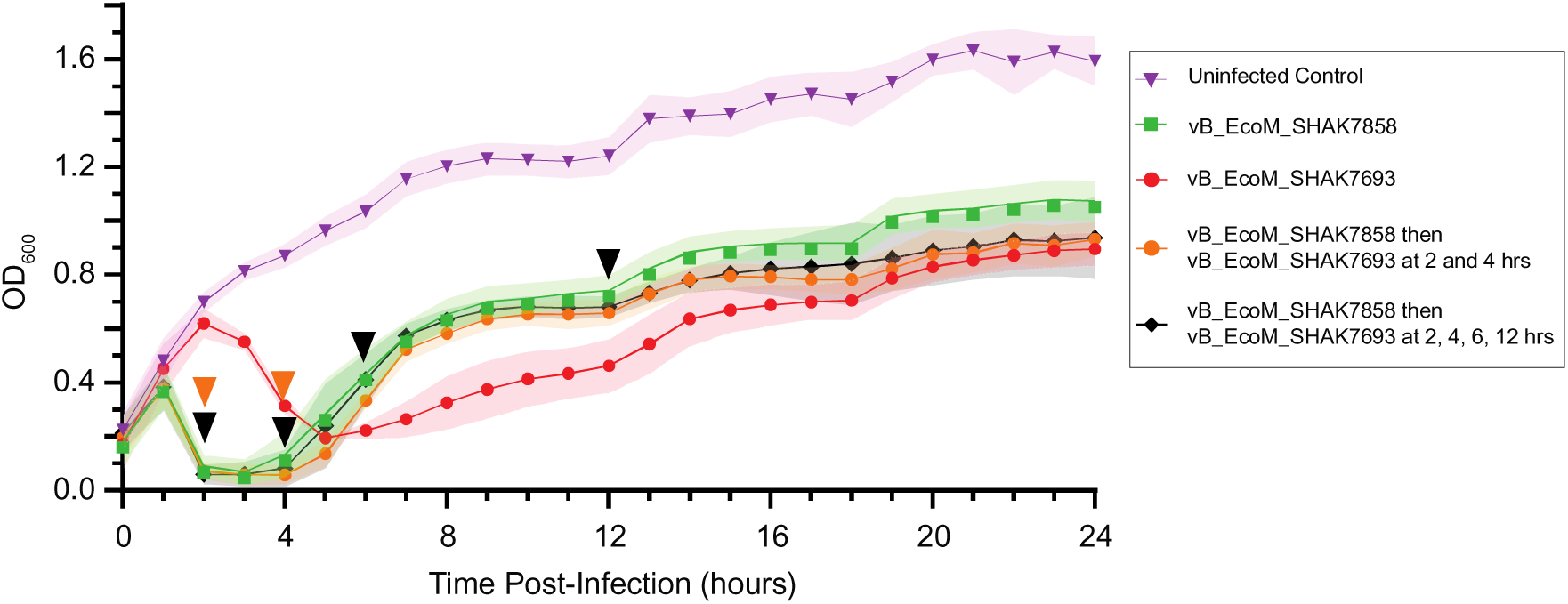
Sequential phage treatment of bacterial strain S112EC with two different phages shows improvement over a single initial dose. Inverted triangles indicate times of sequential dose and colour indicates dosing schedule.

## DISCUSSION

In this work we isolated, sequenced, and annotated six new lytic phage effective against UPEC. For the most part, the genomes of these phages closely conform in architecture and gene content to previously sequenced and characterised phages (Figures S2–S4). *Kagunavirus* vB_EcoM_SHAK7693 had the lowest proportion of CDSs assigned to PHROGs (81.0%) as well as the highest number (49) and proportion (57.6%) of identified CDSs with unknown function (Figure 2), and thus represents the most novel isolated phage in this set.

### Phage host ranges

The similarity of vB_EcoM_SHAK7163 and vB_EcoM_SHAK7704 with Vectrevirus K1E is supported by their ability to efficiently lyse hosts with K1 capsular antigen (PA45B, MS7163, and S79EC) (Table 2). One surprising exception is that vB_EcoM_SHAK7163 has a low efficiency of plating on the K1 antigen strain S79EC (Table 2). One possible explanation is that S79EC has a constellation of predicted antiphage defence systems, distinct from the other K1 strains PA45B and MS7163, including AbiE, AbiP2, Dynamins, Lamassu, Mok Hok Sok, retrons, and PsyrTA (Table S1). These systems mainly defend through an abortive infection mechanism whereby phage-infected bacteria with activated defence systems prevent phage from completing their lifecycle and producing infectious virions. The infected bacteria will accomplish this through induction of programmed cell death or growth arrest (Rousset and Sorek 2023). The overall effect of these abortive infection defence systems is a reduction or elimination of the phage infection in the population (Sberro et al. 2013; Dy et al. 2014; Millman et al. 2020; Millman et al. 2022). By contrast, dynamins appear to work through a different mechanism, where they stabilise the host membrane and delay lysis, thus slowing the phage infection (Guo et al. 2022). These systems may display some differential activity on vB_EcoM_SHAK7163 versus vB_EcoM_SHAK7704 based on one or more of the relatively small differences between these phages, which at the DNA level are 96.3% identical over 94% of their length (Figure 3). Phages vB_EcoM_SHAK7163 and vB_EcoM_SHAK7704 may also have some differential interactions with K5 capsule strains, which are all non-permissive for vB_EcoM_SHAK7163, while two-thirds of the same strains are permissive for vB_EcoM_SHAK7704 (Table 2). The opposite pattern can be seen for the K2 capsule host CFT073 strain for these two phages, which is non-permissive for vB_EcoM_SHAK7704, while vB_EcoM_SHAK7163 can efficiently infect (Table 2). Lastly, vB_EcoM_SHAK7854 and vB_EcoM_SHAK9454 may not be able to use the R1 LPS outer core as a receptor, as they are non-permissive for all R1 strains tested (Table 2).

### Receptor binding domain similarity across different phage families

We observed similar receptor binding domains at the C-terminus of vB_EcoM_SHAK9454 and vB_EcoM_SHAK7858 tail fibers (Figure 4C) corresponding to similar host ranges, while the N-terminus of the proteins was markedly different, likely corresponding to the different requirements for receptor binding, as was seen in the K1-specific phages previously (Stummeyer et al. 2006). Evolutionary receptor binding domain swapping between closely and distantly related phages have been recently shown to govern their specificity towards the O-antigen of hosts (Pas et al. 2023). Phage vB_EcoM_SHAK9454 is an interesting case, as it is very similar to phage T7 (Figure S4) but can efficiently lyse a wide range of UPEC strains (Table 2) across K1, K2, and K5 capsule types, although it seems that R1 LPS strains may be non-permissive (Table 2). This makes vB_EcoM_SHAK9454 a potentially useful chassis for engineering UPEC phage (Ando et al. 2015; Weynberg and Jaschke 2019; Mutalik and Arkin 2022) as the extensive knowledge of phage T7 and its tail fibers, which play important roles in host recognition and DNA delivery in the *Autographiviridae* (Casjens and Molineux 2012; Garcia-Doval and van Raaij 2012), can be used to guide its modification (Molineux 2006; Ando et al. 2015; Huss et al. 2021; Klumpp et al. 2023),.

### Phage cocktail formulation

We experimentally tested cocktails of our isolated phages in several ways: (1) to define lytic efficiency when phage with differential lysis kinetics were used against a single permissive host (Figure 5), (2) to understand how cocktails of phage work against mixtures of permissive and non-permissive hosts (Figure 6), and (3) to compare sequential versus simultaneous dosing schedules (Figure 7). We observed minimal benefit from using two phages (vB_EcoM_SHAK7163 and vB_EcoM_SHAK7693) from different sub-families (*Studiervirinae* and *Guernseyvirinae*) against a single permissive strain (PA45B) compared to only using vB_EcoM_SHAK7163 (Figure 5). This result is in line with previous work, showing a lack of synergistic activity in some phage cocktails, which are not more efficient than a single phage (Pereira et al. 2016).

By contrast, when mixtures of vB_EcoM_SHAK7854 (*Kayfunavirus*) and vB_EcoM_SHAK7693 (*Kagunavirus*) were used against a mixture of S96EC, which was permissive for vB_EcoM_SHAK7854 but non-permissive for vB_EcoM_SHAK7693, and PA45B which was permissive for vB_EcoM_SHAK7693 but non-permissive for vB_EcoM_SHAK7854, we observed robust synergistic activity (Figure 6). Similar effects were seen in recent work of coliphage against mixtures of AMR *E. coli* strains, whereby synergistic effects between phages with differing host ranges were seen (Benala et al. 2023), as well as in other similar studies (Manohar et al. 2019). A sequential dosing study with our two isolated *Kagunavirus* phages (vB_EcoM_SHAK7693 and vB_EcoM_SHAK7858) with high efficiency of plating against the host strain showed that the fast lytic kinetics of vB_EcoM_SHAK7858 could be effectively combined with the more durable suppressive effects of vB_EcoM_SHAK7693 (Figure 7). These results are in line with previous work showing a mild benefit to adding each phage sequentially, although they did observe similar kinetic effects as we did at early time points (Hall et al. 2012).

## Acknowledgements

The authors thank Phil Hugenholtz, Lyman Ngiam, Seweryn Bialasiewicz, Brian Kemish, Leanne Dierens, Soo Jen Low, Paul Evans, Margaret Butler, Helen Pennington and Australian Centre for Ecogenomics (ACE) Sequencing facility for their input and laboratory support.

## Author Contributions

Shahla Asgharzadeh Kangachar (Conceptualization, Data curation, Formal analysis, Investigation, Methodology, Project administration, Software, Validation, Visualization, Writing – original draft, Writing – review & editing), Dominic Y. Logel (Investigation, Methodology, Visualization), Ellina Trofimova (Investigation, Methodology), Hannah X Zhu (Investigation, Methodology), Julian Zaugg (Software, Writing – review & editing), Mark A. Schembri (Resources, Funding Acquisition, Writing – review & editing), Karen D. Weynberg (Conceptualization, Funding Acquisition, Methodology, Project administration, Resources, Validation, Supervision, Writing – review & editing), Paul R. Jaschke (Conceptualization, Formal analysis, Funding acquisition, Investigation, Project administration, Supervision, Visualization, Writing – original draft, Writing – review & editing)

## Data availability

The data used and/or analysed in this study are available from the corresponding author upon reasonable request.

