## Supplementary material for "Discovery and characterisation of new phage targeting uropathogenic *Escherichia coli*": Figure S1, Figure S2, Figure S3, Figure S4, Figure S5, Table S1, Table S2, Table S3

**CONTENTS**

SUPPORTING FIGURES

Figure S1: Large terminase subunit phylogenetic tree

Figure S2. Pairwise CDS comparison of isolated *Autographiviridae* phage with their closest sequence matches.

Figure S3. Pairwise CDS comparison of isolated *Guernseyvirinae* sub-family phage with closest sequence matches.

Figure S4. Pairwise CDS comparison of isolated *Autographiviridae* phage with T7 phage.

Figure S5. Similarities between predicted tail fiber and tail spike structures.

SUPPORTING TABLES

Table S1. Antiphage defence system predictions for UPEC strains used in this study.

Table S2: Phage cocktails against a 1:1 mixture of S96EC and PA45B bacterial hosts.

Table S3: Phage cocktails against S112EC bacterial host.

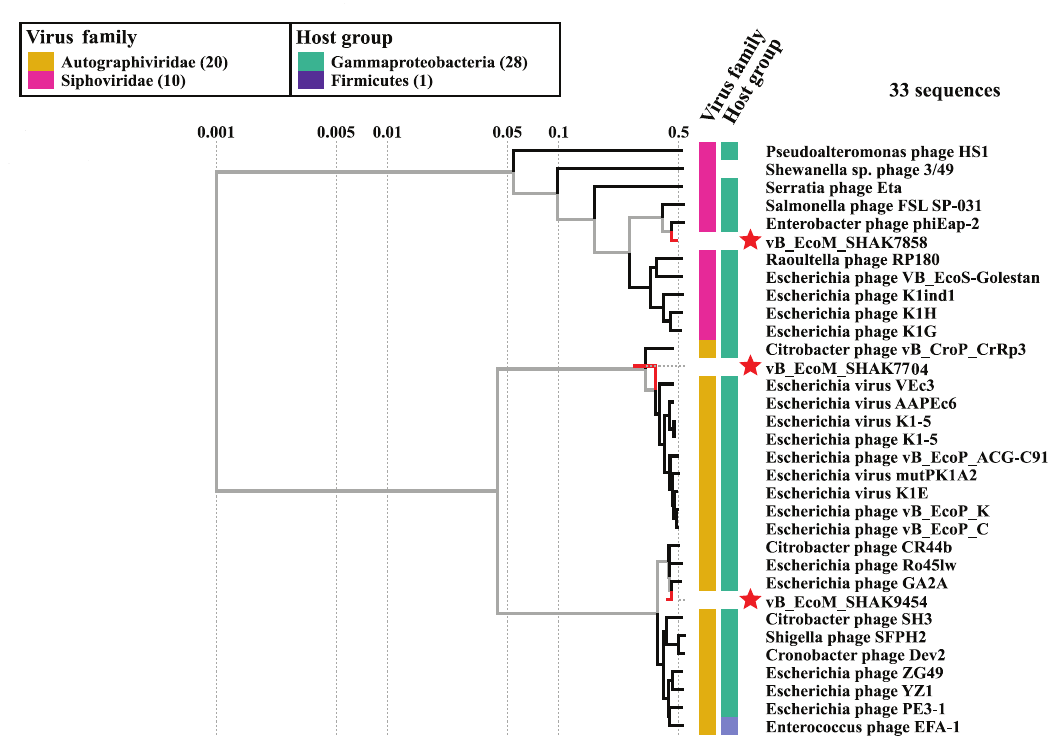

**Figure S1.** Large terminase subunit phylogenetic tree. Three representative phages from this study are highlighted with red stars. The numbers along the top represent bootstrapping values which is a measure of support for the displayed node topology. Generated with VipTree (Version 1.9.1).

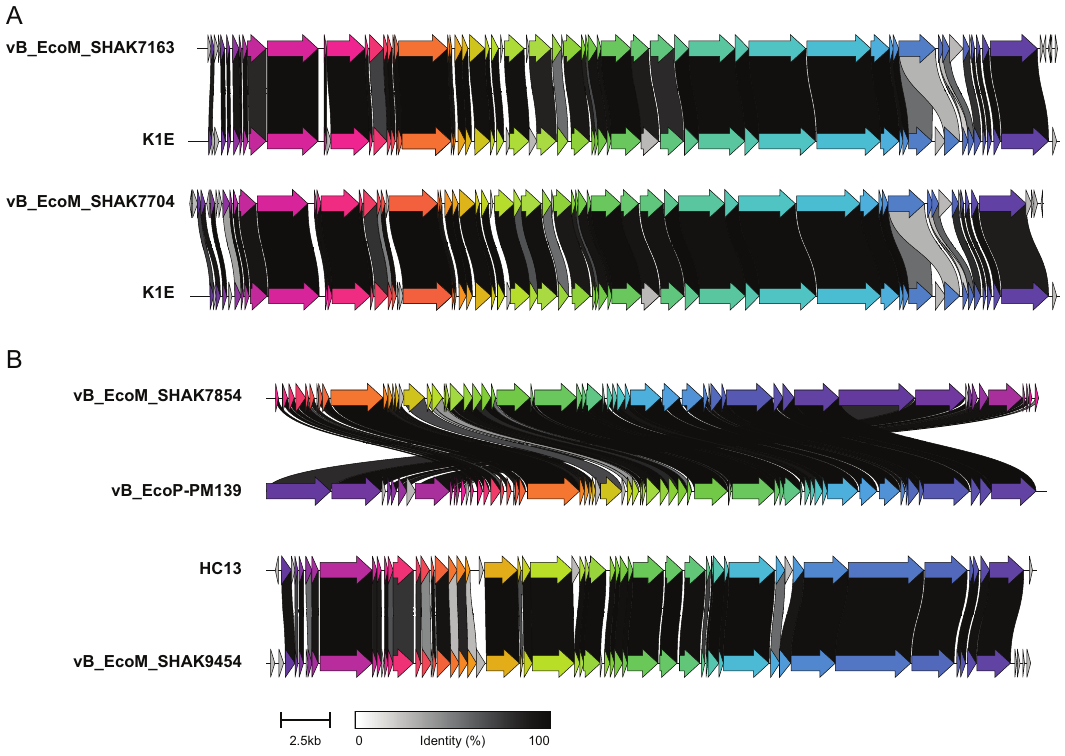

**Figure S2.** Pairwise CDS comparison of isolated *Autographiviridae* phage with their closest sequence matches. (A) *Molineuxvirinae* sub-family phage, (B) *Studiervirinae* sub-family phage. Generated using clinker web interface CAGECAT (the online CompArative GEne Cluster Analysis Toolbox). Homologous CDSs are colored similarly and linked through grayscale bars based on percentage amino acid identity.

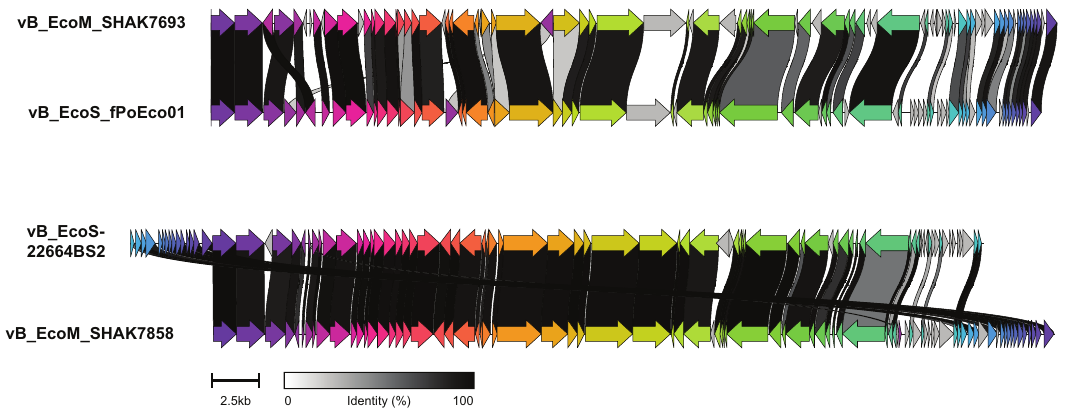

**Figure S3.** Pairwise CDS comparison of isolated *Guernseyvirinae* sub-family phage with closest sequence matches. Generated using clinker web interface CAGECAT (the online CompArative GEne Cluster Analysis Toolbox). Homologous CDSs are colored similarly and linked through grayscale bars based on percentage amino acid identity.

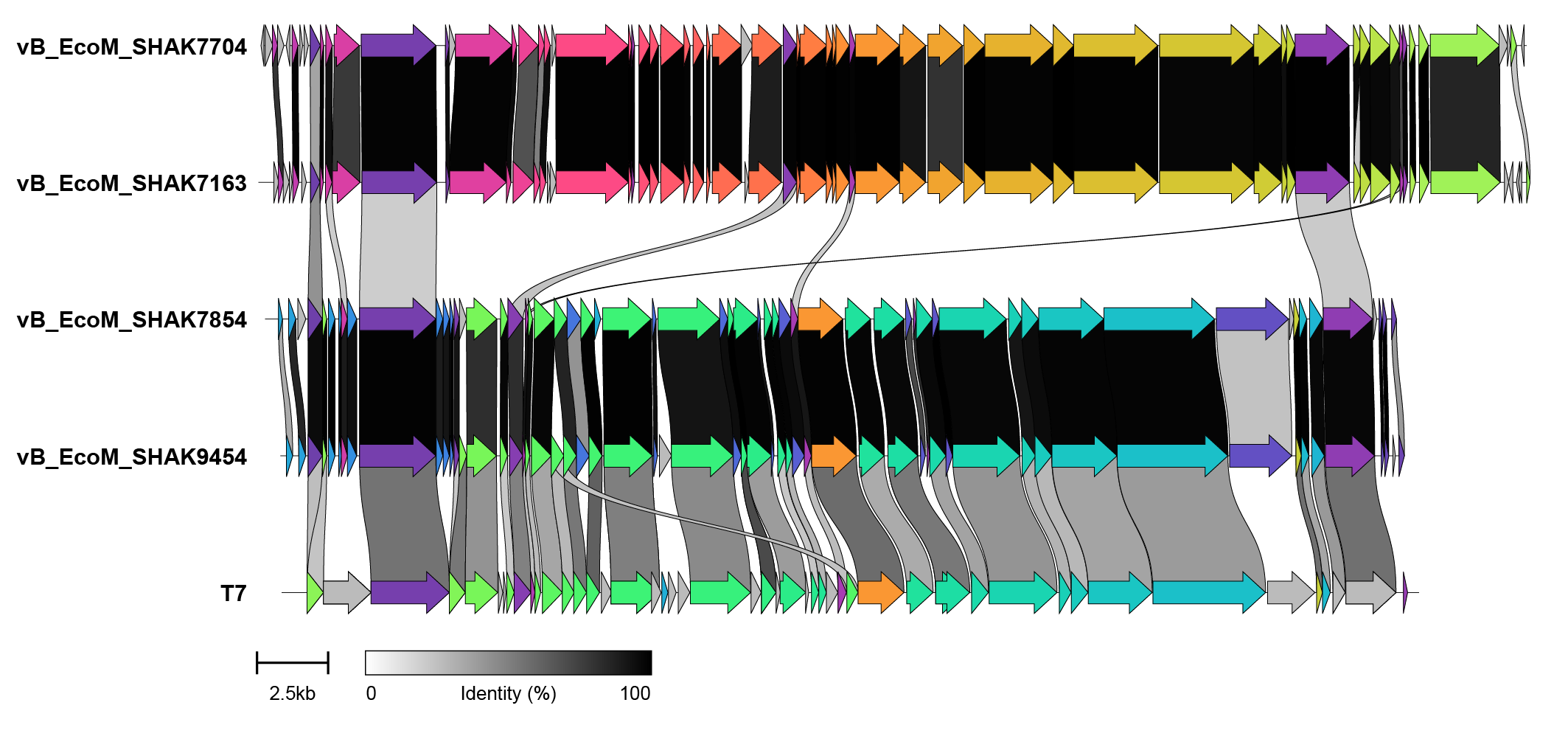

**Figure S4.** Pairwise CDS comparison of isolated *Autographiviridae* phage with T7 phage. Top pair are *Molineuxvirinae* sub-family phage, middle pair are *Studiervirinae* sub-family phage. Generated using clinker web interface CAGECAT (the online CompArative GEne Cluster Analysis Toolbox). Homologous CDSs are colored similarly and linked through grayscale bars based on percentage amino acid identity.

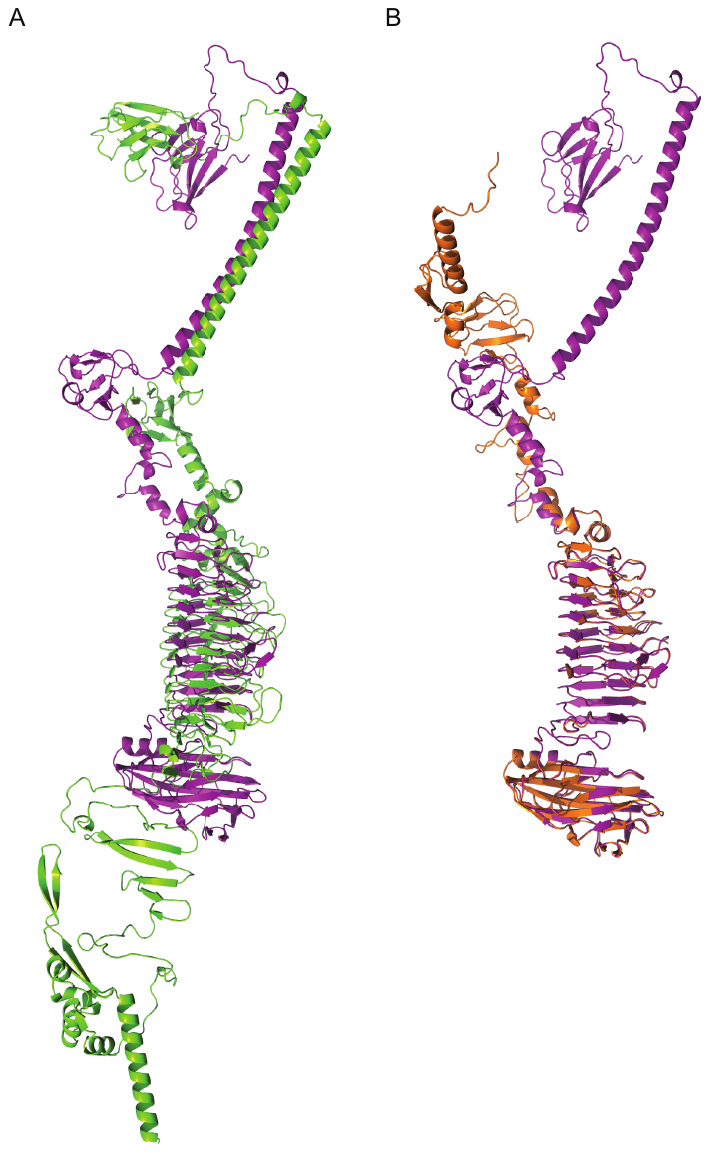

**Figure S5.** Similarities between predicted tail fiber and tail spike structures. (A) Phage vB_EcoM_SHAK7854 (green) tail spike overlapped with vB_EcoM_SHAK9454 (purple) tail fiber. (B) Phage vB_EcoM_SHAK7858 (orange) tail fiber overlapped with vB_EcoM_SHAK9454 (purple) tail fiber. AlphaFold predicted structures for each protein were aligned using PyMOL.

**Table S1.** Antiphage defence system predictions for UPEC strains used in this study.

| **Strain** | **AbiE** | **AbiH** | **AbiP2** | **AbiU** | **BREX (type I)** | **Cas type I-E** | **Cas type I-F1** | **CRISPR array** | **CBASS type I** | **DMS_other** | **DRT class I** | **DRT class II** | **Druantia type I** | **Druantia type III** | **Dsr1** | **dXTPase** | **dCTPdeaminase** | **Dynamins** | **Gabija** | **Gao19** | **Gao29** | **Gao Ape** | **Gao Qat** | **Gao RL** | **Gao Tmn** | **Hma** | **Lamassu** | **Mok Hok Sok** | **Mokosh type II** | **PD-Lambda-1** | **PD-T4-1** | **PD-T4-3** | **PD-T4-7** | **PD-T7-1** | **PD-T7-5** | **PsyrTA** | **Retron_I-C** | **RM type I** | **RM type II** | **RM type III** | **RM type IV** | **SanaTA** | **Septu type I** | **SoFIC** | **Thoeris type I** | **Thoeris type II** | **Total** |
| --- | --- | --- | --- | --- | --- | --- | --- | --- | --- | --- | --- | --- | --- | --- | --- | --- | --- | --- | --- | --- | --- | --- | --- | --- | --- | --- | --- | --- | --- | --- | --- | --- | --- | --- | --- | --- | --- | --- | --- | --- | --- | --- | --- | --- | --- | --- | --- |
| **PA45B** |  |  |  |  |  |  | 1 | 1 | 1 | 1 |  |  |  |  |  |  |  |  |  | 1 |  | 1 | 1 |  |  |  |  |  | 1 | 1 |  | 1 |  |  |  |  |  |  | 1 |  | 1 |  |  | 1 | 1 | 1 | 14 |
| **S96EC** | 1 |  |  |  |  |  |  |  | 1 | 1 |  |  |  |  |  |  |  |  |  |  |  |  |  |  |  |  | 1 | 1 | 1 |  |  |  |  |  |  | 1 | 1 | 1 | 1 |  | 1 |  |  |  |  |  | 11 |
| **S112EC** | 1 |  |  |  |  |  |  |  | 1 | 1 |  |  |  |  |  |  |  |  |  |  |  |  |  |  |  |  | 1 | 1 | 1 |  |  |  |  |  |  | 1 | 1 | 1 | 1 |  | 1 |  |  |  |  |  | 11 |
| **MS7163** |  |  |  |  |  |  | 1 | 1 | 1 | 1 |  |  |  |  |  |  |  |  |  |  |  |  | 1 |  |  |  |  |  | 1 |  |  | 1 |  | 1 |  |  |  | 1 | 1 |  | 1 |  |  |  |  |  | 10 |
| **S39EC** | 1 |  |  |  | 1 |  |  |  |  | 1 |  |  |  |  |  |  |  |  | 1 |  |  |  |  |  | 1 |  | 1 | 1 | 1 |  |  |  |  |  |  | 1 | 1 | 1 | 1 |  | 1 |  |  |  |  |  | 13 |
| **CFT073** | 1 |  |  |  |  |  |  |  | 1 | 1 |  |  |  |  |  |  |  |  |  |  |  |  | 1 |  |  |  |  |  |  |  |  |  |  |  |  | 1 |  | 1 | 1 | 1 |  |  |  |  | 1 | 1 | 10 |
| **PA7B** |  |  |  |  |  |  |  |  |  | 1 |  |  |  |  |  | 1 |  |  |  |  |  |  |  |  |  |  | 1 | 1 |  |  |  |  |  | 1 |  | 1 |  | 1 | 1 | 1 | 1 |  | 1 |  | 1 | 1 | 13 |
| **S101EC** | 1 |  |  |  | 1 |  |  |  |  | 1 |  |  |  |  |  |  |  |  | 1 |  |  |  |  |  | 1 |  | 1 | 1 | 1 |  |  |  |  |  |  | 1 | 1 | 1 | 1 |  | 1 |  |  |  |  |  | 13 |
| **S79EC** | 1 |  | 1 |  |  |  |  |  |  | 1 | 1 |  |  |  |  |  |  | 1 |  |  |  |  |  |  |  |  | 1 | 1 |  |  |  |  |  |  |  | 1 | 1 | 1 | 1 |  | 1 |  |  |  |  |  | 12 |
| **S129EC** | 1 |  |  |  | 1 |  |  |  | 1 | 1 |  | 1 |  |  |  |  | 1 |  |  |  |  |  |  |  |  |  | 1 | 1 | 1 |  |  |  |  |  |  | 1 | 1 | 1 | 1 |  | 1 | 1 |  |  |  |  | 15 |
| **S116EC** | 1 |  |  |  |  |  |  |  | 1 | 1 |  |  |  |  |  |  |  |  |  |  |  |  |  |  |  |  | 1 | 1 | 1 |  |  |  |  |  |  | 1 | 1 | 1 | 1 |  | 1 |  |  |  |  |  | 11 |
| **HVM204** | 1 |  |  |  |  |  |  |  |  | 1 |  |  |  |  | 1 |  | 1 |  |  |  | 1 |  |  | 1 |  |  | 1 | 1 | 1 |  |  |  |  |  |  | 1 | 1 | 1 | 1 |  | 1 |  |  |  |  |  | 14 |

Predictions based on a combination of PADLOC, DefenseFinder, and CRISPRDetect using default parameters.

**Table S2.** Phage cocktails against a 1:1 mixture of S96EC and PA45B bacterial hosts.

| **Condition** | **Phages Combined in Cocktail** | | | |
| --- | --- | --- | --- | --- |
| 1 | 9454 | 7854 | - | - |
| 2 | 9454 | 7163 | - | - |
| 3 | 9454 | 7693 | - | - |
| 4 | 7854 | 7163 | - | - |
| 5 | 7854 | 7693 | - | - |
| 6 | 7163 | 7693 | - | - |
| 7 | 9454 | 7854 | 7163 | - |
| 8 | 9454 | 7854 | 7693 | - |
| 9 | 9454 | 7163 | 7693 | - |
| 10 | 7854 | 7163 | 7693 | - |
| 11 | 9454 | 7854 | 7163 | 7693 |

**Table S3.** Phage cocktails against S112EC bacterial host.

|  | **Time (hrs)** | | | | | |
| --- | --- | --- | --- | --- | --- | --- |
| **Treatment #** | **0** | **2** | **4** | **6** | **12** | **18** |
| 1 | 7858 | 7858 |  |  |  |  |
| 2 | 7858 | 7693 |  |  |  |  |
| 3 | 7858 | 7693+ |  |  |  |  |
| 4 | 7858 | 7858 | 7858 |  |  |  |
| 5 | 7858 | 7693 | 7693 |  |  |  |
| 6 | 7858 | 7693+ | 7693+ |  |  |  |
| 7 | 7858 |  | 7858 |  |  |  |
| 8 | 7858 |  | 7693 |  |  |  |
| 9 | 7858 |  | 7693+ |  |  |  |
| 10 | 7858 | 7858 | 7858 | 7858 |  |  |
| 11 | 7858 | 7693 | 7693 | 7693 |  |  |
| 12 | 7858 | 7693+ | 7693+ | 7693+ |  |  |
| 13 | 7858 | 7858 | 7858 | 7858 | 7858 |  |
| 14 | 7858 | 7693 | 7693 | 7693 | 7693 |  |
| 15 | 7858 | 7693+ | 7693+ | 7693+ | 7693+ |  |
| 16 | 7858 |  |  |  | 7858 |  |
| 17 | 7858 |  |  |  | 7693 |  |
| 18 | 7858 |  |  |  | 7693+ |  |
| 19 | 7858 | 7858 | 7858 | 7858 | 7858 | 7858 |
| 20 | 7858 | 7693 | 7693 | 7693 | 7693 | 7693 |
| 21 | 7858 | 7693+ | 7693+ | 7693+ | 7693+ | 7693+ |

| 7858 | vB_EcoM_SHAK7858_(MOI=10) |
| --- | --- |
| 7693 | vB_EcoM_SHAK7693_(MOI=10) |
| 7693+ | vB_EcoM_SHAK7693 (MOI=100) |
